# Divergence Model of Protein Evolution

**DOI:** 10.1101/045930

**Authors:** Darrell O. Ricke

## Abstract

Analysis of the evolutionary conservation of amino acids is useful for the analysis of sequence variants detected in individuals with regard to possible impact on protein function. Rapid advances in DNA sequencing technologies are enabling affordable access to SNPs, exome sequencing, and whole genome shotgun sequencing. An understanding of the biological processes that have shaped diverging protein sequences aids the interpretation of sequence variants. The divergence model of protein evolution is presented as a general framework for interpreting information available from comparative analysis of protein sequences.

## Introduction

Protein missense mutations are either neutral, deleterious to protein function, or confer a selective advantage (rare). Over time adaptive mutations are selected for and become fixed within a population and deleterious mutations are selected against[1]. Neutral mutations drift through populations[2]. To assess the functional importance of protein amino acid residues, sequences from multiple species are aligned in a multiple sequence alignment. When available, protein structures provide insights into the structural roles and importance of residues[3, 4].

Analysis of aligned sequences led to the molecular clock proposal [5], matrix tables of protein evolution mutation frequencies[6], predictions of gene evolutionary divergence history depicted as phylogenetic trees [7], observations of invariant and highly changeable positions [8], and observation of pressure against radical interior changes coupled with considerable less pressure against surface residues [9]. [10] observed that insertions and deletions tend to be small (<= 5 amino acid residues) and occur in regions of loops and random coils. Bottema et al. [11] proposed that proteins are essentially composed of two types of amino acids: critical (essential for protein function) and spacer (serving as peptide backbone spacers for proper structural folding of critical residues). Sequence alignments are used to obtain new insights into protein function, structure, and disease relevance.

The evolutionary record observed with sequence comparisons is shaped by the rates and patterns of mutations. Different relative rates of mutations are observed with transitions more frequent than transversions and with even lower rates for insertions and deletions[12-14]. Organisms that use 5-methyl-cytosine have a 5^th^ DNA base with a substantially higher mutation rate than non-methylated cytosine residues[15, 16]. Evolutionary rates of mutation are coupled to the DNA repair fidelity with mitochondrial DNA rates of mutations higher than nuclear rates. The consistency of DNA repair rates enables the phylogenetic estimates from the molecular clock proposal.

Dickerson [9] observed that genes evolve at different rates. Genes on the X chromosome appear to accumulate mutations at a slightly lower rate than the autosomes[17]. Highly expressed genes appear to accumulate fewer mutations because of transcription-coupled repair[18]. Polak and Arndt[19] also observed transcription induced mutations at the 5’ end of human genes. Observed differences between mutations at silent residues compared to rates of accumulation of missense mutations are interpreted as evidence of adaptation. Evolutionary models do not accommodate more than simple patterns of mutation or the concept that some residues are essential for function and missense mutations at these positions are deleterious. The divergence model of protein evolution is presented that models positive, negative, and neutral mutations in the context of essential and nonessential residues as a general framework for understanding residue importance and how protein sequences are diverging.

## Methods

### Divergence Model of Evolution

Taking into account the evolutionary clock proposal, neutral theory, critical spacer model, adaptation, and the patterns of observed mutations, it is possible to model the evolutionary divergence of protein sequences as a stochastic process within the functional constraints of the protein. The observed missense mutations (M) observed over millions of years of evolutionary time (N) is modeled as the sum of accumulated mutations (*p*_*i*_ + *q*_*i*_)^*N*_*i*_^ – *q*_*i*_^*N*_*i*_^ = (1 – *q*_*i*_^*N*_*i*_^) applied to variable residues accounting for adaptation (*α*_*i*_*R*) and fast mutating 5-methyl-cytosines (1 – *s*_*i*_^*N*_*i*_^)β_*i*_*R*:

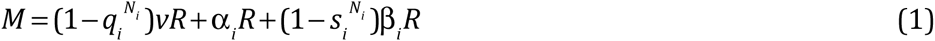

Where

- M - number of observed missense mutations
- i - species or divergence segment
- *p*_*i*_ - rate of mutation in 1 million years for species i
- *q*_*i*_ = 1 – *p*_*i*_ - the probability of a residue not being mutated in 1 million years for species i
- *N*_*i*_ - number of millions of years of divergence time
- *v* - fraction of variable residues in a protein
- *R* - total number of residues in a protein
- *α*_*i*_ - fraction of residues with adaptation mutations for species i
- *r*_*i*_ - rate of mutation at residues with 5-methyl-cytoside in 1 million years for species i
- *s*_*i*_ = 1 – *r*_*i*_ - the probability of a 5-methyl-cytoside residue not being mutated in 1 million years for species i
- β_*i*_ - the fraction of residues with 5-methyl-cytoside bases where mutations will cause nonsynomous mutations
- γ - the fraction of critical or invariant residues for the protein with 1 = *v* + α_*i*_ + β_*i*_ + γ

The invariant residues in a multiple sequence alignment can be used as an approximation for the γ*R* residues with inclusion of some variant residues with no observed mutations (*q*^*N*^*vR*). From an alignment, the α_*i*_*R* residues can be approximated from conserved alignment positions by comparing different taxonomic classes (e.g., birds versus mammals)[20]. A similar example for hemoglobin subunit beta is shown in Figure S1. Likewise, β_*i*_*R* residues can be roughly approximated from estimating ancestral residues with 5-methyl-cytosine bases in codon bases 1 or 2 for Arginine CGN and Proline CCG codons: R/H/C and R/W/Q, and L/P alignment positions. Other CG-dinucleotides are possible but are less easily identified from protein alignments. Note that Ricke *et al*.[21] observed L/P segments in signal peptides. Being able to exclude these alignment residue positions from consideration, it is possible to reduce equation 1 to the simplified equation 2.

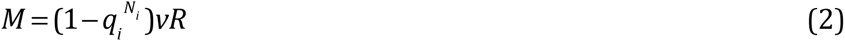

For two sequences being compared, if they experienced approximately the same rates of mutations, equation 2 can be simplified to equation 3 with V representing the remaining alignment positions excluding extended regions lacking conserved residues to avoid errors introduced from misalignment of residues. From the multiple sequence alignment positions V, it is possible to estimate q and N for observed mismatches M.

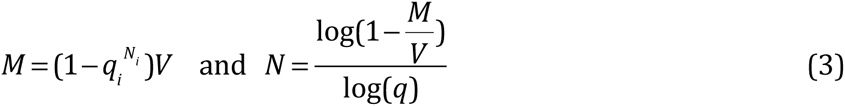

### Approximating Ancestral Residues with 5-Methyl-Cytosine

CpG-dinucleotides represent 1/16 of the possible dinucleotides. For Eukaryotes, not all CpG-dinucleotides are methylated. In Table 1, evidence for CpG-dinucleotides with 0, 1 and 2 methylated bases are apparent based on observed mutations at CGN arginine residues in TP53[22]. The methylation status of CpG-dinucleotides can be estimated by the observed residue changes in a multiple sequence alignment. Arginine (CGN codons) residues with methylated CpG-dinucleotides will exhibit missense mutations with a higher probability for transition mutations (see Table 1). Other codons with potential methylated CpG-dinucleotides include Proline (CCG), Serine (TCG), Threonine (ACG), Alanine (GCG), Valine (GTN), Asparagine (GAY), Glutamine (GAR), and Glycine (GGN). To avoid confusion with non-5-methyl-Cytosine mutations, only transition mutations at Arginine (CGN) codons and Proline (CCG->CTG) to Leucine transition mutation patterns in a multiple sequence alignment will be considered as candidates for fast mutating 5-methyl-cytosine mutations to approximate the βR residues.

**Table 1.** Estimating CpG-dinucleotide Methylation Patterns in sample TP53 Arginine CGN residues[22]

| Codon | Residue | Transitions | Transversions | Methylated Bases |
| --- | --- | --- | --- | --- |
| CGC | R156 | H10, L3 | G3, C3, P26, S3 | 0 |
| CGC | R158 | H74, L63 | C17, G11, P10 | 0 |
| CGC | R175 | H904 | L20, C16, G16 | 1 |
| CGC | R181 | H24 | C19, P12 | 0 |
| CGA | R196 | Q4, *170 | P17, L1 | 1 |
| CGA | R213 | Q29, *258 | L39, P6, G2 | 1 |
| CGC | R248 | Q617, W522 | P17, L73, G14 | 2 |
| CGG | R267 | Q12, W29 | P17, L7, G1 | 0 |
| CGT | R273 | H567, C524 | P29, S14, L96, G10 | 2 |
| CGG | R282 | Q25, W439 | P17, G29, L3 | 1 |
| CGC | R283 | H12 | P24, C17, L4, G2 | 0 |

### Model Evaluation

FASTA sequences were extracted from the SwissProt[23] dat file with the Java SwissProtParser[24] program. This program annotates sequences with metadata including gene name, taxonomy, organism name, etc. in the FASTA description field. FASTA sequences were partitioned by gene with the Ruby split_by_gene.rb program. The split_by_gene program uses the gene name annotation to create files for each gene. Genes were analyzed with the Ruby diverge.rb program. The divergence program creates a multiple sequence alignment of the protein sequences and characterizes the aligned sequences using the divergence model. To avoid including class-specific adaptation mutations, variable residues are estimated only from Eutheria sequences. Alignment positions consistent with likely ancestral 5-methyl-cytosine bases were identified and excluded. Extended variable regions with more than 14 spacers are excluded to avoid potential regions of uncertain alignments. Variable residues for Eutheria are estimated using both the average number of observed mismatches at variable residues with N_Eutheria_ set to 100 million years. The Dendroscope[Huson *et al*. 2007] program was used to visualize newick files generated by the diverge.rb program.

## Discussion

Vertebrate sequences from the SwissProt database were extracted and genes analyzed with the divergence model. The fraction of variable residues *vR* and the probability of a residue not being mutated, *q*, were estimated (when possible) for each gene. The estimated values for *vR* and *q* by gene are included in supplemental table S1. Proteins with smaller lengths and low *v* values provide fewer alignment positions for calculations and estimates to be based on. The density of variable to essential residues within a protein can vary by functional domain and should be taken into consideration for partial protein sequences. Calculated *v* and *q* values are shown in Figures 2 and 3 for 1,344 SwissProt vertebrate genes from Table S1. Examples of N estimates are shown in Figure S2 for TP53 and Figure S3 for hemoglobin subunit beta. The widely accepted dogma that genes evolve at different rates appears to really be primarily that genes have different proportions of essential versus unessential residues. When the evolution rate does very considerably, then there is likely relevant biology responsible or possibly data errors.

**Figure 1.**
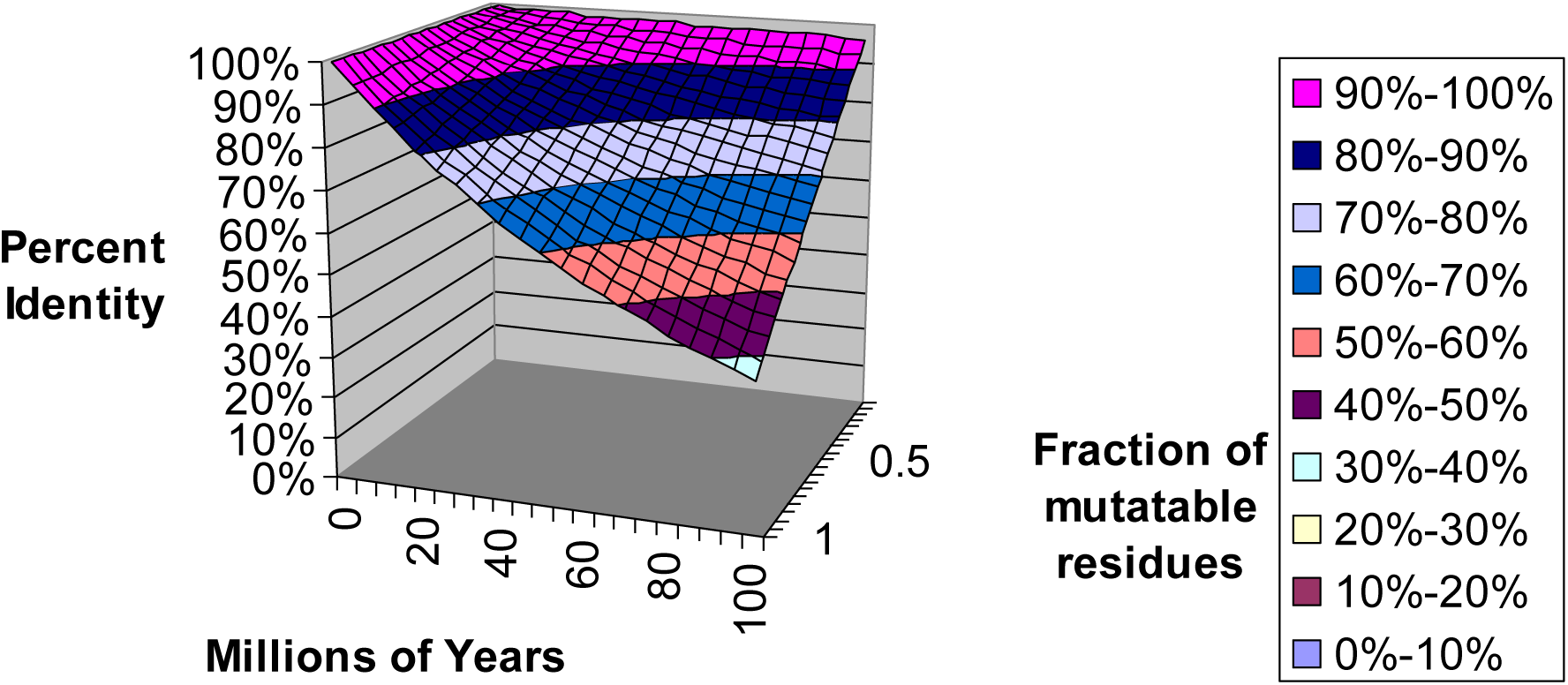
Theoretical Divergence by gene for v and N. Estimated sequence divergence from equation 3 for divergence time (N) and the fraction of variable residues (v) in a protein.

**Figure 2.**
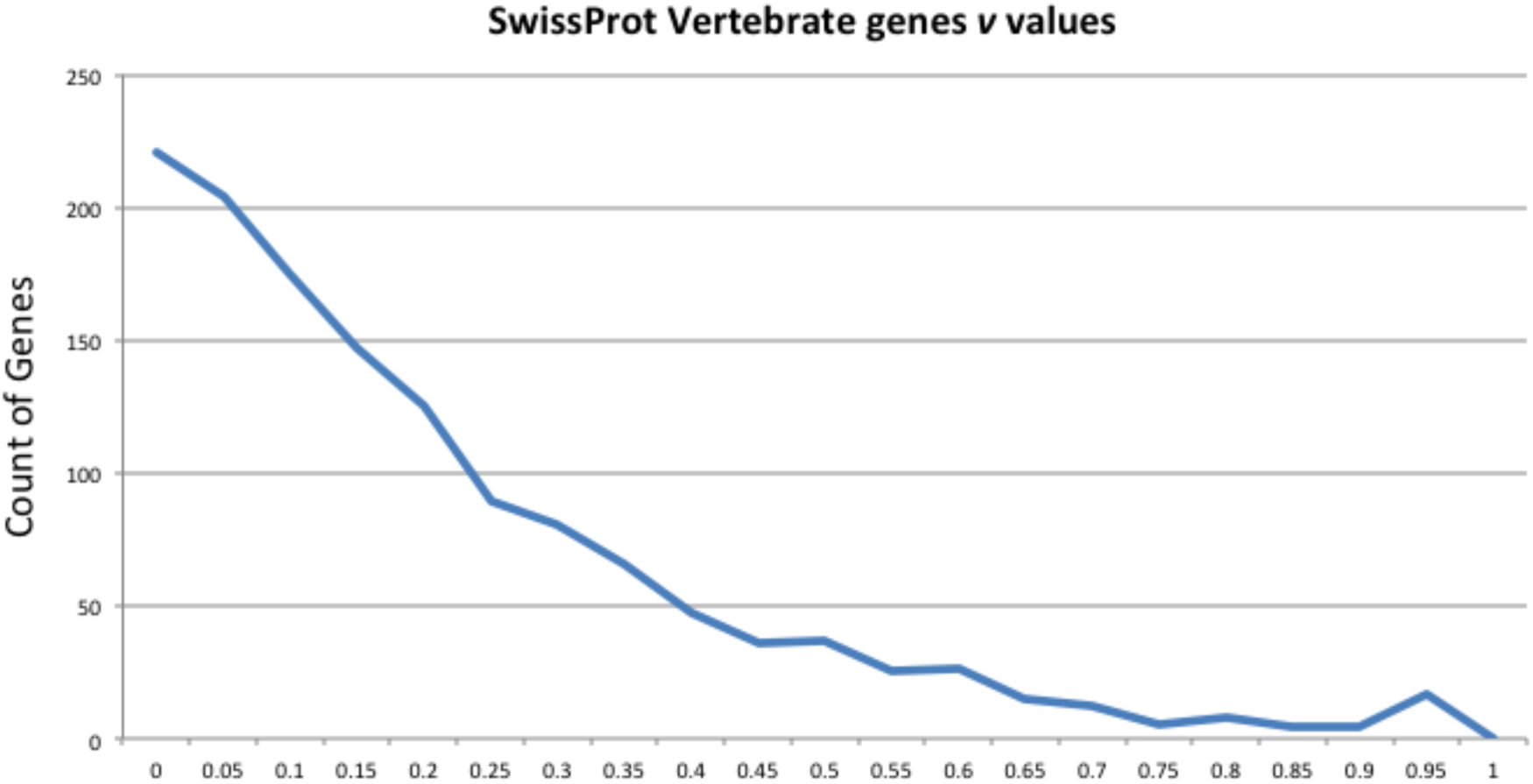
Vertebrate genes v values.Distribution of 1,344 genes from SwissProt vertebrates by v value.

**Figure 3.**
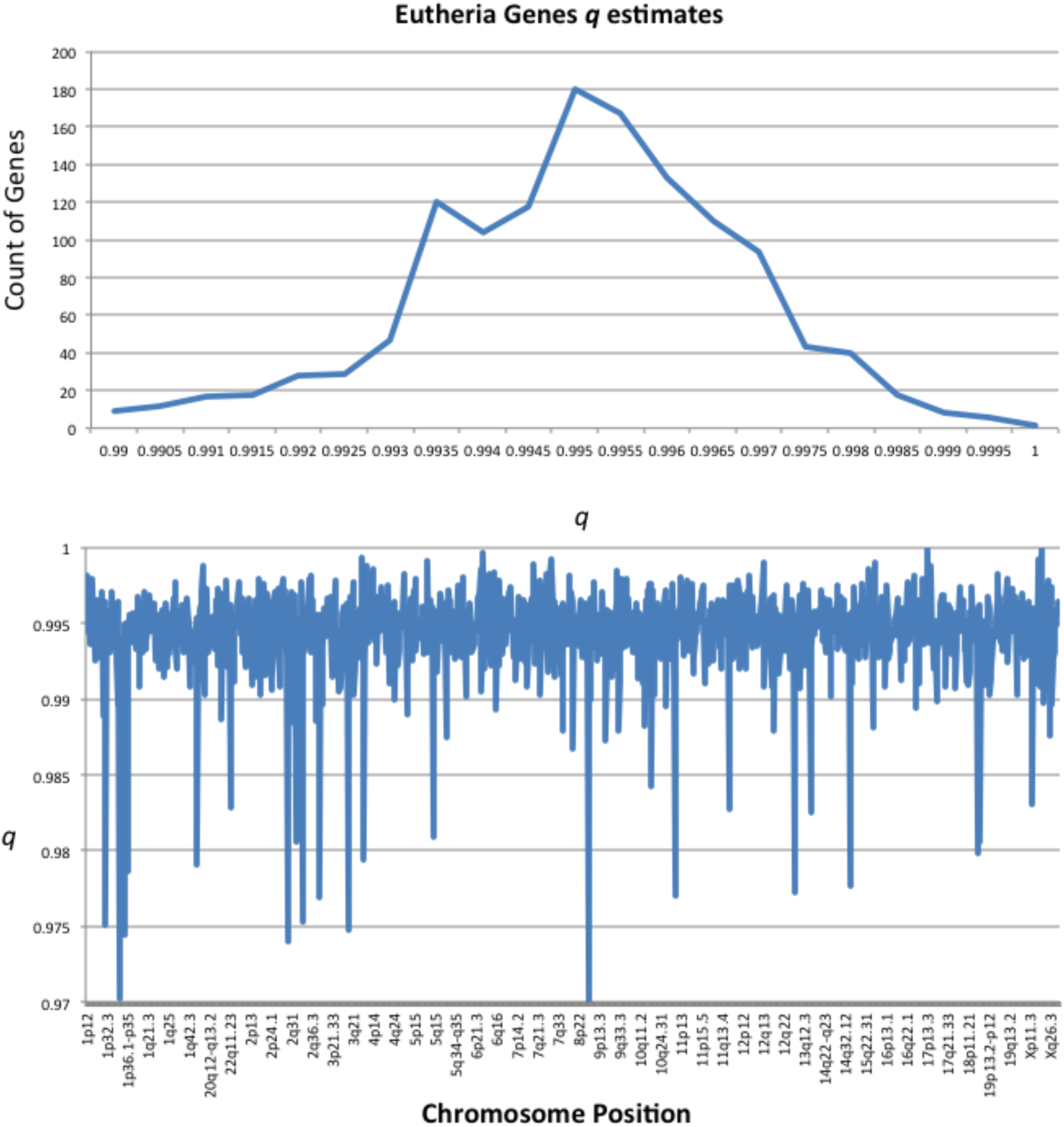
Eutheria estimates of q values, (a) Distribution of 1,344 genes by q value; (b) Distribution of q by human chromosome position.

### Observations

1. The apparent molecular evolutionary clock is a direct result of the fidelity of cellular DNA repair processes combined with stochastic mutations with characteristic patterns of mutations.

a. 5-methyl-cytosine residues mutate at a much higher rate than other residues and have a higher probability of being observed for closely related species. On average, these represent roughly 1/32, or less, of the variable residues being considered for a gene.
2. The ratio of critical to variable residues for a gene is essentially stable across evolutionary time with small variations due to insertions and deletions.
3. Adaptation mutations can be observed in multiple sequence alignments and should be taken into consideration when classifying a residue as conserved or non-conserved.
4. Orthologs with extended path lengths exceeding stochastic random variability compared to orthologs with comparable diverge time are candidates for functioning close to paralogs than orthologs. Likewise, orthologs with too small of N are candidates for horizontal gene transfers, species misidentification errors, etc.
5. Estimates of N need to take into consideration the stochastic nature of mutations in the context of protein function and adaptations and only provide a general relative estimate of divergence time. Estimated values for N are sensitive to estimated q value.

## Conclusions

The Divergence model of protein evolution is presented as a general model for understanding the divergence of protein sequences from common ancestral sequences. The model accommodates positive, negative, and neutral mutations in the context of protein function. The model provides the framework for interpreting information from comparative sequence analysis. An implementation of this model is available on GitHub [https://github.com/doricke/Divergence].

## Supplemental Figure Legends

Figure S1. Hemoglobin beta multiple sequence alignment.

Figure S2. TP53 N estimates visualization with Dendroscope.

Figure S3. Hemoglobin beta N estimates visualized with Dendroscope.

